# The contribution of plant life and growth forms to global gradients of vascular plant diversity

**DOI:** 10.1101/2023.03.06.531444

**Authors:** Amanda Taylor, Patrick Weigelt, Pierre Denelle, Lirong Cai, Holger Kreft

## Abstract

Plant life and growth forms (shortened to ‘plant forms’) represent key functional strategies of plants in relation to their environment and provide important insights into the ecological constraints acting on the distribution of biodiversity. Despite their ecological importance, how the spectra of plant forms contribute to global gradients of plant diversity is unresolved.

Using a novel dataset comprising >295,000 species, we quantify the contribution of different plant forms to global gradients of vascular plant diversity. Further, we establish how plant form distributions in different biogeographical regions are associated with contemporary and paleoclimate conditions, environmental heterogeneity, and phylogeny.

We find a major shift in representation by woody perennials in tropical latitudes to herb-dominated floras in temperate and boreal regions, following a sharp latitudinal gradient in plant form diversity from the tropics to the poles. We also find significant functional differences between regions, mirroring life and growth form responses to environmental conditions, which is mostly explained by contemporary climate (18-87%), and phylogeny (6-62%), with paleoclimate and heterogeneity playing only a minor role (<23%).

This research highlights variation in the importance of different plant forms to diversity gradients worldwide, providing a much-needed quantification for long-standing ideas and concepts structuring plant assemblages.

## Introduction

Plant life and growth forms (shortened to ‘plant forms’) are non-phylogenetic classifications of species that share similar trait combinations (e.g., herbs, geophytes, lianas), and represent key ecological strategies of plants in relation to their environment. While the grouping of taxa into plant forms has a long-standing history in ecology (Humboldt, 1806; Warming & Knoblauch, 1895; Raunkiaer, 1934; Box, 1981; Ellenberg & Mueller-Dombois, 1982), the concept is still widely used in modern-day studies on functional diversity (Mayfield *et al.*, 2005; Leroy *et al.*, 2021), biome and vegetation classifications (Box & Fujiwara, 2015; Mucina, 2019), ecological restoration (Gómez-Aparicio, 2009), and plant systematics (Govaerts *et al.*, 2021). Despite this long period of development and being a major facet of plant life-history variation, how the spectra of plant life and growth forms contribute to global biodiversity patterns remains unquantified. For instance, while terrestrial herbs are characteristic of temperate grasslands, and trees of tropical rainforests, exactly how much they contribute to total plant diversity in or outside of these biomes, and in relation to coexisting plant forms, are questions that have yet to be addressed on a global scale due to a lack of data that became available only recently.

Indeed, several plant forms may coexist within a single region, yet may differ substantially in their proportional representation. Trees offer an exceptional example, as while globally diverse in tropical forests (Cazzolla Gatti *et al.*, 2022), their regional contribution to tropical biodiversity is comparable to co-occurring terrestrial herbs, epiphytes, and climbers (Spicer *et al.*, 2020). Similarly, plant forms within distinct biomes may comprise species that originate from different evolutionary lineages in different biogeographical regions, leading to variation in their contribution within equivalent biomes worldwide. For instance, the large Bromeliaceae family endemic to the New World contributes significantly to the epiphyte flora in Neotropical forests, explaining to some degree the comparatively poorer epiphyte diversity in tropical forest biomes in other botanical realms (Taylor *et al.*, 2022). Another case study is the surprisingly poor representation of native stem succulents in Australia, despite the subcontinent sharing many climatic affinities with the trans-continental succulent biome, which has favoured the establishment of several invasive succulent species (Ringelberg *et al.*, 2020). Thus, regions that share similar climatic conditions may not necessarily share the same functional composition, highlighting the importance of historical biogeography, and the need to consider the legacy of past climate on contemporary plant diversity patterns.

Sudden fluctuations in climate can promote species extinctions, particularly of poorer dispersers that have limited ability to migrate, and whose ranges are less likely to intersect with climate refugia (Sandel *et al.*, 2011). In other cases, species physiologies may prevent the evolution of adaptations to tolerate increasingly unstable climate conditions. The low diversity of epiphytes in temperate forests, for instance, has been partly related to the distribution of ice cover during the Last Glacial Maximum (Taylor *et al.*, 2022), possibly leading to a reduction in the ‘epiphyte niche’ (Zotz, 2005), and that epiphytes are likely poorly adapted to frost (but see Zotz and Hietz, 2001. On the contrary, geophytes, which tend to favour seasonally dry climates, have been shown to decrease in species richness with increasing historical climate stability (Sosa & Loera, 2017), despite climate stability being a major driver of biodiversity by promoting speciation (Fine, 2015). Understanding how past climate changes have influenced present day plant form distributions can ultimately help to predict future climate-driven shifts in vegetation structure and functioning.

Evolutionary history may also play an important role in governing the contribution of different plant forms to biodiversity. Under the same environmental conditions, phylogenetically related species may be ecologically similar in terms of life history, resource use, interaction partners, and consequently, functional traits (Burns & Strauss, 2011). On the other hand, distantly related lineages may independently evolve similar traits in different biogeographical realms, as has been shown for C4 photosynthesis (Sage *et al.*, 2012), and other specialised metabolisms (Pichersky & Lewinsohn, 2011), insular woodiness (Zizka *et al.*, 2022), shrubby stature (Gehrke *et al.*, 2016), and stem succulence (Ringelberg *et al.*, 2020). It is also not uncommon for members of the same plant form to have drastically different phylogenetic constraints, such as the strong phylogenetic constraints on tropical evergreen tree lineages that do not have perennating buds and different wood anatomy compared to deciduous trees (Zanne *et al.*, 2014). The variable contribution of plant forms to biodiversity patterns could therefore elucidate how the complex life history dimensions of plants are modified by the environment and provide important new insights into the evolutionary and ecological constraints underpinning plant trait- and diversity-environment relationships.

Here, we provide a global assessment of the importance of different plant forms to global patterns of vascular plant diversity (see Engemann *et al.*, 2016 for the New World), and address how their relative contributions in different biogeographical regions are associated with contemporary and paleoclimate conditions, environmental heterogeneity, and phylogeny. We consider two of the most widely accepted and basic classifications of plant forms - ‘life forms’ and ‘growth forms’, which for centuries have been used to study vegetation around the world (Humboldt, 1806; Warming & Knoblauch, 1895; Raunkiaer, 1934; Box, 1981; Ellenberg & Mueller-Dombois, 1982). The term ‘growth form’ refers to the growth habit of plants, recognising trees, shrubs, subshrubs, terrestrial herbs, climbers, and epiphytes as distinct functional types (defined in Table 1). Life forms, in contrast, describe the different biological tolerances and strategies of plants to withstand unfavourable seasonal environmental conditions based on the location of perennating buds (Raunkiaer, 1934; Ellenberg & Mueller-Dombois, 1982). Therophytes, hemicryptophytes, geophytes, chamaephytes, and phanerophytes (Table 1) have vastly different survival strategies, ranging from persisting as seeds (therophytes) and underground storage organs (geophytes), to having perennating buds at (hemicryptophytes), near (chamaephytes), or well above the soil surface (phanerophytes). Accordingly, therophytes and geophytes can persist in arid and cold environments, hemicryptophytes and chamaephytes in cool-temperate climates, and many phanerophyte species are restricted to regions without frequent frost or drought (Sosa & Loera, 2017; Howard *et al.*, 2019; Irl *et al.*, 2020; Taylor *et al.*, 2021). While several more advanced classification schemes exist today (e.g., Halloy, 1990), we argue that because life and growth forms serve as a baseline classification of plant assemblages worldwide, they represent some of the most accessible and documented plant traits to date.

**Table 1.**
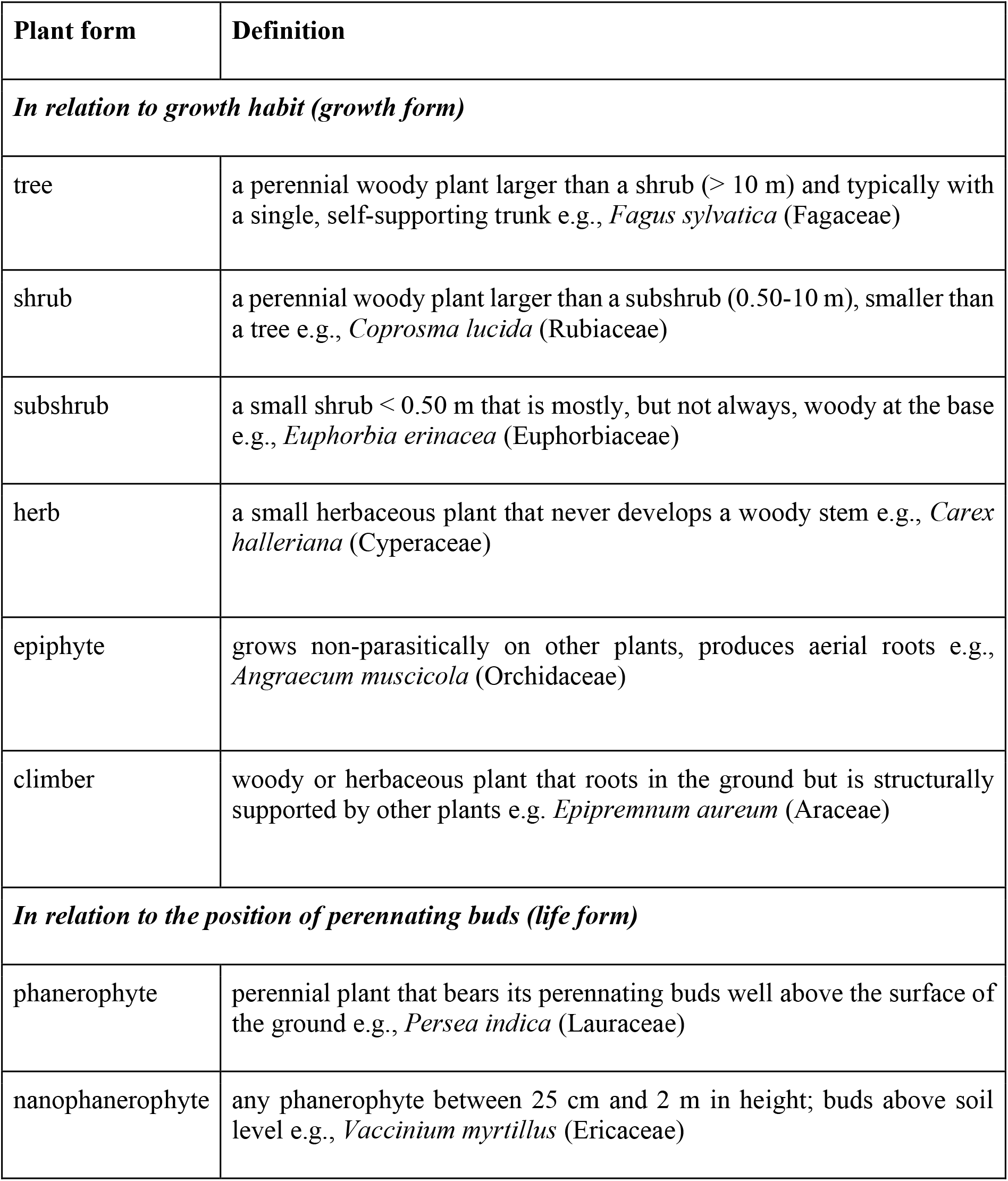
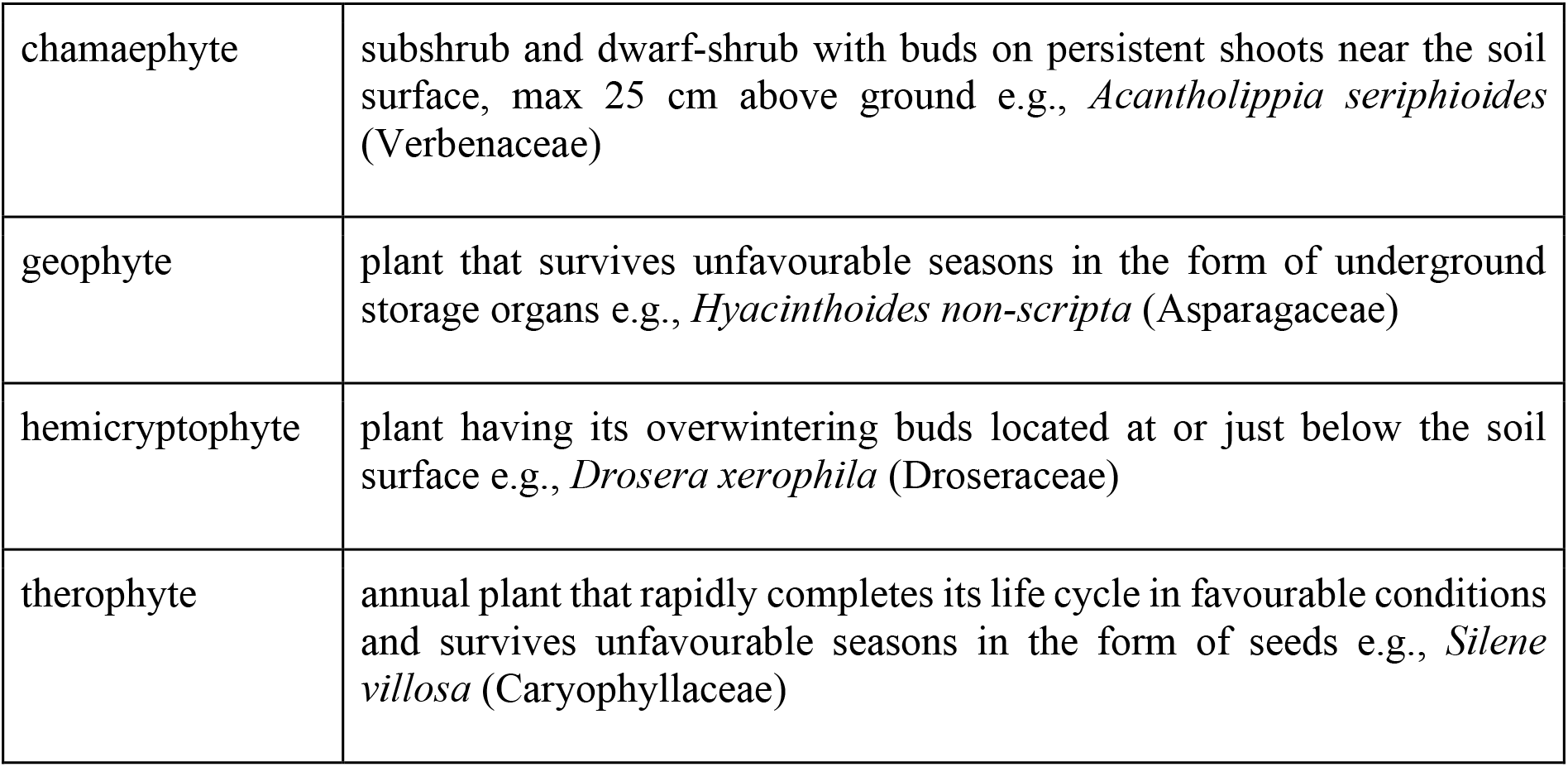
Major life and growth forms defined by plant growth habit (growth form) and the position of perennating buds (life form). Although epiphytes were originally classified as phanerophytes within the Raunkiaer life form scheme (Raunkiaer, 1934), they were later separated due to their unique arboreal habit (Ellenberg & Mueller-Dombois, 1982). As their revised classification is not related to the position of perennating buds, we include them here as a growth form, rather than a life form.

Using a global dataset consisting of over 295,000 vascular plant species, we quantify the relative contribution of life and growth forms to biodiversity in different regions around the world. As several studies have shown that plant diversity and the distribution of biomes and vegetation types can be broadly predicted by gradients of temperature and precipitation (Whittaker, 1975), climate heterogeneity (Reu *et al.*, 2011), and past climate changes (e.g., palms, Blach-Overgaard *et al.*, 2013), we expect that similar climatic variables will be important predictors of plant form distributions. We broadly expect that (*H_1_*) the proportional representation of woody plants will decrease from the tropics to the poles and be inversely related to the proportional representation of terrestrial herbs. We also expect that (*H_2_*) variation in the composition and contribution of plant forms in different regions around the world will be driven primarily by gradients of precipitation, temperature, past climate change, heterogeneity, and phylogeny, the effects of which will depend on individual plant form adaptations.

## Material and Methods

### Biodiversity data

We merged two of the world’s largest botanical databases - the World Checklist of Vascular Plants (WCVP version 2.0, Govaerts *et al.*, 2021) and the Global Inventory of Floras and Traits (GIFT version 3.0, Weigelt *et al.*, 2020) to obtain a near-complete global checklist of life and growth form classifications. Both databases provide comprehensive global checklists of over 344,000 accepted species that come from a variety of resources, including published taxonomies, floras, and regional checklists. Additionally, these databases are now freely available to the public, with the WCVP being accessible here (https://powo.science.kew.org/ under the tab “DATA”), and GIFT here (https://github.com/BioGeoMacro/GIFT).

We obtained from the WCVP and GIFT databases plant form classifications for 223,338 and 250,599 species, respectively, with most growth forms coming from GIFT, and life forms from WCVP. Due to epiphytes often being incorrectly classified, we did not derive epiphyte classifications from either database, but instead from a recently published epiphyte list (Zotz *et al.*, 2021), which amounted to 24,279 species, totalling 295,755 species once all databases were merged. All species classified as epiphytes from the epiphyte list were removed as other plant forms to reduce biases in the representation by non-trees (i.e., an epiphyte cannot simultaneously be an ‘epiphyte’ and ‘herb’). Thus, all ‘herbs’ only include terrestrial herbs. However, in other cases where multiple classifications were assigned to a single species, we took all possible combinations to account for some variability in plant forms exhibited by individual species across different environments (i.e., an herb in one region may appear shrubby in another region). Together, our dataset accounts for 87% of all accepted plant species, with the missing 23% being due to a lack of plant life or growth form classification, or due to having a classification outside of our 12 major plant forms, which we did not consider due to small sample sizes. For example, we did not consider hydrophytes due to consisting of just ~870 species. Species names across all databases and lists follow the WCVP taxonomic backbone.

Merging these databases provided several benefits beyond a comprehensive global dataset of plant form classifications. For one, species distributions from the WCVP are recorded at the level of ‘Botanical Country’ of the World Geographical Scheme for Recording Plant Distributions (WGSRPD, Brummitt et al., 2001). GIFT, however, offers checklists at finer resolutions in some places, with species distributions also being aggregated at the regional (e.g., departments or states) to country-level (e.g., Peru) in other places. To obtain the highest resolution of plant form distributions, we always opted to use checklists from the smaller regions provided that 1) a complete checklist of native vascular plants was available, and 2) the smaller regions were perfectly nested within larger regions, where if not we would take the larger geographical entity (e.g., if we were missing two departments of Ecuador, we would not use the remaining departments, but instead aggregate species to the entire country of Ecuador). To this end, 56% of the regions included in this study come exclusively from GIFT, and 64% from WCVP and GIFT combined, out of a total of 564 continental mainland and island regions (Table S3). We excluded oceanic islands because of their insularity and unique biogeographic history, which can lead to deviations in plant functional traits (e.g., Taylor *et al.*, 2019).

Because some plant species have more than one life or growth form assignment, we aggregated the number of species belonging to each plant form to obtain total ‘growth form’ and ‘life form’ richness, which was then used to calculate the relative proportion of each plant form within a region. More specifically, while ‘absolute species richness’ is the sum of all *species* within a region, ‘growth form richness’ and ‘life form richness’ is the sum of all species of a particular growth or life form within a region, respectively. Each plant form was then divided by its respective total richness to obtain its relative contribution to biodiversity within each region (e.g., number of trees/sum of (trees + shrubs + subshrubs + herbs + climbers + epiphytes)). To determine how the contribution and composition of plant forms varies among large biogeographical regions, we followed the same procedure for each major botanical continent; Africa, Australia, North America, South America, Asia Temperate, and Asia Tropical, as well as the dominant biome (in terms of total area of occupancy) of each geographical entity (e.g., Namibia = “Deserts & Xeric Shrublands”, Cyprus = “Mediterranean Forests, Woodlands & Scrub”). We excluded from this analysis biomes that were dominant in only one region to reduce region-specific biases. As such, we removed the ‘mangrove’ and ‘rock and ice’ biomes due to them being dominant in single regions, Guinea-Bissau, South and Greenland, respectively.

To gain a general overview of the diversity of plant forms around the world, we calculated the Shannon diversity index (*H*’), which considers both the number of species within each plant form in a given region and their relative abundance (measured here as proportional representation). We then took the exponential of this index to quantify the Shannon effective number of plant forms, which measures the number of equally abundant/common plant forms needed to produce a particular observed value of *H* (Jost, 2006).

### Phylogenetic data

To account for phylogenetic relatedness among species, and to tease apart the evolutionary and ecological contributions of plant forms to global richness patterns, we calculated phylogenetic eigenvectors at the region level and included them as additional predictor variables in our global models. We used the extended version of the Smith and Brown (2018) phylogeny, implemented in the R package ‘V.PhyloMaker2’ (Jin & Qian, 2022), choosing the World Plants GBOTB.extended.WP.tre extension due to our taxonomic nomenclature being most similar to this database (https://www.worldplants.de, Hassler, 2000). We then bound missing species to a respective congener before executing the phylo.maker function to make a phylogenetic tree hypothesis under scenario 3 (Jin & Qian, 2022). Although scenario 3 adds missing species as polytomies, several studies have shown the robustness of this approach when used in large-scale, biogeographical analyses (Qian & Jin, 2016, 2021; Cai *et al.*, 2022).

In a second step, we calculated phylogenetic turnover between regions separately for each life and growth form, opting for the Simpson index given its insensitivity to variation in species richness among sites (Baselga, 2010). However, similar in the way that phylogenetic richness is strongly influenced by species richness, species compositional turnover should also have a profound influence on phylogenetic turnover. We therefore quantified the standardised effect size of phylogenetic turnover (Phylo_ses_) for each plant form, correcting for variation in compositional turnover among regions using the ‘phyloregion’ package in R (Daru *et al.*, 2020).

This was done by shuffling the tip names of each plant form-subsetted phylogeny 1,000 times to create multiple null assemblages and taking the difference between the observed phylogenetic turnover (Phylo_obs_) and the mean null phylogenetic turnover (Phylomean_null), divided by the standard deviation (Phylo_sd_); Phylo_ses_ = (Phylo_obs_ - Phylo_mean_null_) / Phylo_sd_. Finally, we ran a Principal Coordinates Analysis (PCoA) on the resulting Phylo_ses_ distance matrix to create phylogenetic eigenvectors at the regional level, which represents the total variation in phylogenetic composition among regions. Used as explanatory variables, these eigenvectors can be used to infer how phylogenetic relatedness of species across regions influences the contribution of plant forms to richness patterns within a given region. We included in our models the first 4 PCoA axes, as together these could explain > 90% of the variation in phylogenetic relatedness among regions.

### Environmental predictor variables

We initially selected 15 candidate predictor variables representing contemporary climate, paleoclimate, and environmental heterogeneity. To assess how life and growth forms are associated with contemporary climate, we selected three metrics derived from CHELSA V.2.1 (30 arc-seconds, Karger *et al.*, 2017), which in previous studies have been shown to play an important role in determining plant diversity patterns (Scheiner & Rey-Benayas, 1994; Kreft & Jetz, 2007; Cai *et al.*, 2022): mean annual temperature (°C), mean annual precipitation (mm), and temperature seasonality (coefficient of variation of mean annual precipitation). These predictors should capture life and growth form strategies that may or may not favour seasonal climates (e.g., temperate zone), regions with extreme temperatures (e.g., deserts, alpine environments), and regions that experience prolonged periods of drought.

To test for potential legacies of historical climate on present day plant form contributions we considered three paleoclimate variables: climate change velocity in precipitation and temperature since the Last Glacial Maximum (m/a, 30 arc-seconds, Hijmans et al., 2005), and climate stability (Owens & Guralnick, 2019). Rapid past climate changes are likely to favour plant forms that can swiftly recover following a disturbance (e.g., some herbs, Haukioja & Koricheva, 2000), while stable climates should favour most plant forms, but especially those restricted in their climate niche (e.g., epiphytes, Taylor *et al.*, 2022). Finally, climate heterogeneity measured as the Euclidean distance to a multivariate climate centroid (reprojected from 50 x 50 km^2^ grids, Fournier *et al.*, 2020) was included to tease apart plant forms that are limited to stable, homogeneous climates (e.g., tropics), from those able to persist in heterogeneous, non-stable climates (e.g., temperate regions).

Several historical climate variables (during the last interglacial and Pleistocene warming period) were considered - mean annual temperature, temperature seasonality, mean annual precipitation, and precipitation seasonality, yet were excluded due to strong correlations (Pearson cor = > 0.70) with contemporary climate variables. Length of growing season, aridity, number of frost days, and actual evapotranspiration were also excluded due to similar issues of collinearity. In the case of correlated variables, we always selected those that were most strongly related to life and growth form representation. One exception is mean annual temperature and temperature seasonality, which despite being correlated (Pearson cor = 0.71) we decided to retain both predictors given that seasonality may strongly reflect plant form strategies to survive harsh conditions, and that temperature is one of the most important drivers of plant diversity (Whittaker, 1975). We also found a higher model performance with their combined inclusion (based on lower AIC values of models with both predictors vs models with temperature and seasonality included separately). A summary of the retained predictor variables used in this study and their hypothesised effect on each plant form is provided in supporting information (Table S1, S2). Further, we attach as an additional supporting file Table S3 our aggregated data sheet with all response and predictor variables used for the analyses.

### Statistical Analysis

Generalised linear mixed effects models (GLMMs) with a binomial error term were used to determine the ecological conditions favouring each plant form, whilst simultaneously controlling for phylogenetic non-independence of species within each region. We included all contemporary climatic and past environmental predictors described above as fixed effects, and botanical continent as a random effect to account for large-scale geographic variation in evolutionary history among regions (Gerstner *et al.*, 2014). Relative proportions of plant forms were included as composite response variables (no. of species in plant form x, number of species in other plant forms).

To test for issues of spatial autocorrelation and over-dispersion, which commonly lead to the inflation of type 1 errors in macroecological analyses (Dormann *et al.*, 2007; Bolker *et al.*, 2009), we ran non-spatial GLMMs as described above, and computed the Moran’s *I* coefficient on our model residuals. As we found only low to moderate spatial dependency, ranging from Moran’s *I =* 0.11, p < 0.05 for hemicryptophytes to Moran’s *I =* 0.27, p < 0.05 for chamaephytes, we decided not to run spatial models. Conversely, we encountered significant over-dispersion across all plant form models (chisq/rdf = > 500, p < 0.05), which was resolved by including an additional observation-level random effect. We considered our final and most parsimonious models as those with the lowest AICc scores and delta AICc = <2 (Burnham & Anderson, 2004). Finally, we calculated the relative importance of contemporary climate, paleoclimate, environmental heterogeneity, and phylogeny on the proportional contribution of life and growth forms to global biodiversity patterns. All statistical analyses and maps were made in R version 4.2.1 with major input from the packages ‘lme4’ (Bates, 2010), ‘jtools’ (Long, 2019), ‘MuMin’ (Barton, 2015), ‘V.Phylomaker2’ (Jin & Qian, 2022), ‘phytools’ (Revell, 2012), ‘phyloregion’ (Daru *et al.*, 2020), ‘lwgeom’ (Pebesma, 2018), and ‘ggplot2’ (Wickham & Chang, 2016).

## Results

### Contribution of life and growth forms to botanical continents and biomes

We found striking differences in the contribution of different plant forms to global biodiversity patterns and in relation to key global drivers of vegetation. Relative growth form composition consisted of 16% trees, 28% shrubs, 12% subshrubs, 32% herbs, 7% epiphytes, and 5% climbers (rounded to the nearest whole number). The composition of life forms consisted of 21% phanerophytes, 20% nanophanerophytes, 15% chamaephytes, 24% hemicryptophytes, 11% geophytes, and 9% therophytes. It is important to remember that some species have >1 plant form classification, and that these proportions are in relation to total growth form richness (n = 325,538) and life form richness (n = 245,129), not species richness.

Terrestrial herbs (incl. herbs, hemicryptophytes, geophytes, therophytes) were most prevalent in the northern botanical continents, accounting for up to 77% of all vascular plants in North America, Europe, and Asia-temperate (Fig. 1a). The contribution by perennial woody plants (tree, shrub, subshrub), in contrast, peaked in South America, Africa, Asia-tropical, and Australasia, comprising between 54% and 63% of the vascular flora. Epiphytes and climbers were generally poorly represented in all regions, except for South America and Asia-tropical, where epiphytes characterised 12% and 13% of the total flora, and climbers 6%, and 8%, respectively.

**Figure 1.**
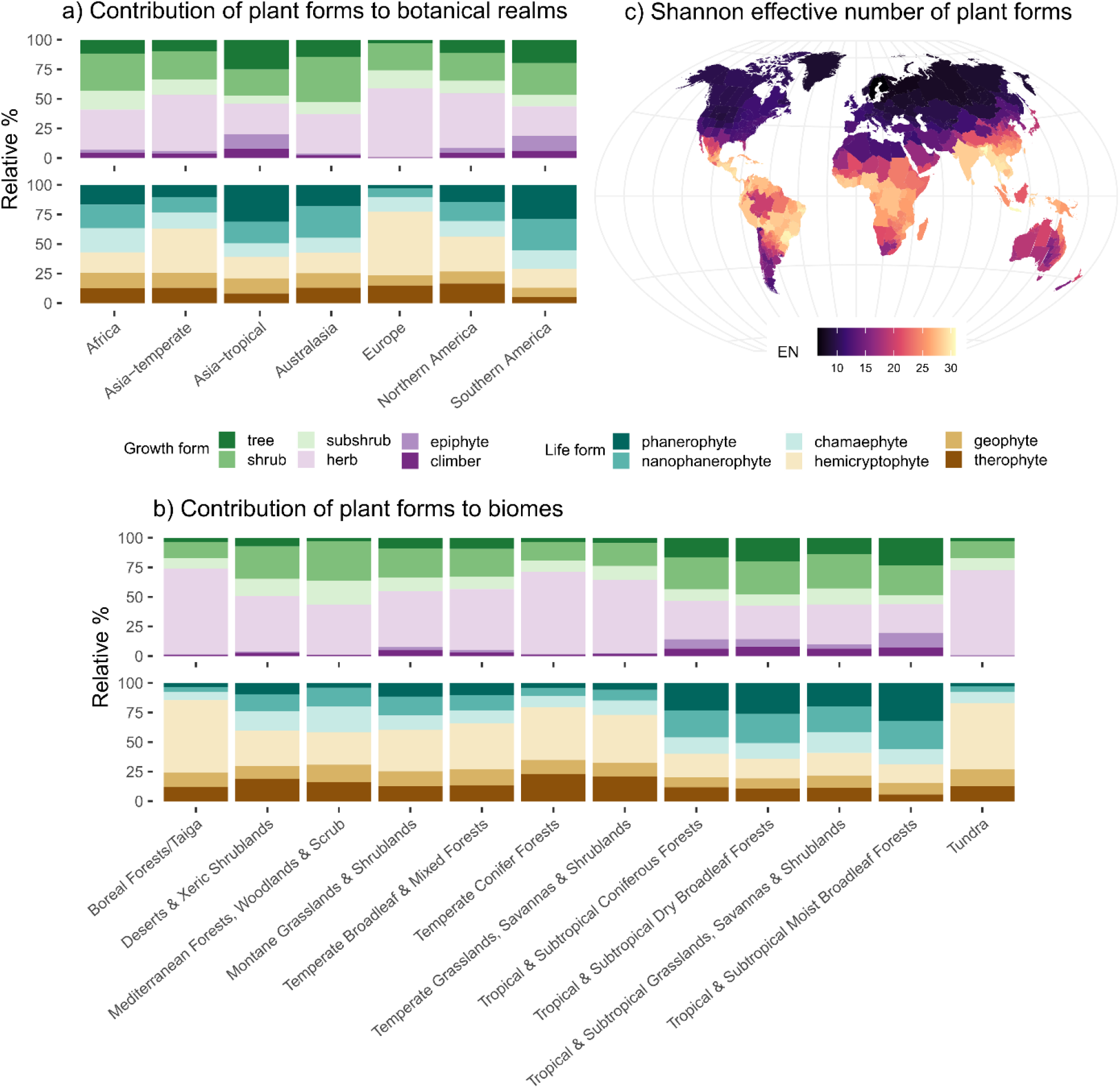
The proportional contribution of life and growth forms to (a) botanical continents and (b) biomes. Figure (c) depicts the global distribution of plant form diversity, quantified here as the effective number of life and growth forms, taken in this case as the exponential of the Shannon–Wiener Index, H’. Map follows the Winkel tripel projection (ESRI:54042).

Comparisons among biomes also revealed variation in the distribution of plant form composition. Species composition of biomes characterised by extreme cold (‘boreal forests/taiga’, ‘temperate conifer forests’, ‘tundra’, ‘montane grasslands and shrublands’) was dominated by herbaceous plant forms, which accounted for up to 85% of the flora. Shrub and subshrub species followed in second place, comprising between 22% and 36% of the flora in these cooler regions. Arid and hot biomes (‘deserts and xeric shrublands’, ‘Mediterranean forests, woodlands and scrub’) were composed mainly of herbs (43-47%), shrubs (27-34%), and hemicryptophytes (27-30%), but also showed notable contributions from subshrubs (15-20%), nanophanerophytes (14-16%), chamaephytes (17-22%), geophytes (11-15%), and therophytes (16-19%).

Changes in the distribution and composition of life and growth forms was evident in the tropical and subtropical biomes (tropical and subtropical: ‘coniferous forests’, ‘dry broadleaf forests’, ‘grasslands, savannas and shrubland’, ‘moist broadleaf forests’), which favoured perennial woody vegetation – trees (14-23%) and shrubs (25-29%), as well as phanerophytes (20-32%) and nanophanerophytes (22-25%), although herbs and hemicryptophytes were also well-represented (24-34% and 16-20%, respectively). The proportional representation of epiphytes (12% in moist broadleaf forests) and climbers (8% in dry broadleaf forests) also peaked in these regions. Finally, ‘temperate broadleaf and mixed forests’ displayed a varied composition of relatively even proportions of trees, subshrubs, chamaephytes, geophytes, nanophanerophytes, phanerophytes, and therophytes (9-14%), yet maintained high proportions of herbs, shrubs, and hemicryptophytes (24-51%, Fig. 1b). Despite variation in the contribution of life and growth forms to plant diversity among the major botanical continents and biomes, we found a clear latitudinal gradient in the diversity of plant forms from the tropics to the poles (Fig. 1c).

### Growth forms

In tropical forests, especially in Southeast Asia and the Neotropics, trees account for the majority of plant diversity (up to 42%, Fig. 2), with shrubs generally being the second most prevailing growth form (up to 24%, see Table S3 for all proportions by region). This is in stark contrast to temperate forests where most plant species are herbs (up to 79% in Europe), while trees generally constituted only a small fraction (<3%) in these same regions. Our models mostly associate this clear variation to gradients of temperature, seasonality, and precipitation, with trees being strongly negatively related to seasonality (standardised estimate Est. = −0.41, p = <0.01), and positively related to precipitation (Est. = 0.38, p = <0.01) and temperature (Est. = 0.35, p = <0.01, Fig. 4a). Herbs and subshrubs displayed an inverse relationship, being positively related to seasonality, and negatively associated to temperature, with subshrubs also being negatively associated to precipitation (Table S4). Arid Mediterranean climates and regions overlaying large desert biomes were composed primarily of shrubs (up to 50%, Fig. 2), subshrubs (up to 27%), and herbs (up to 75%), which was driven by varying combinations of temperature, seasonality, precipitation, and phylogeny (Table S4, Fig. 4a).

**Figure 2.**
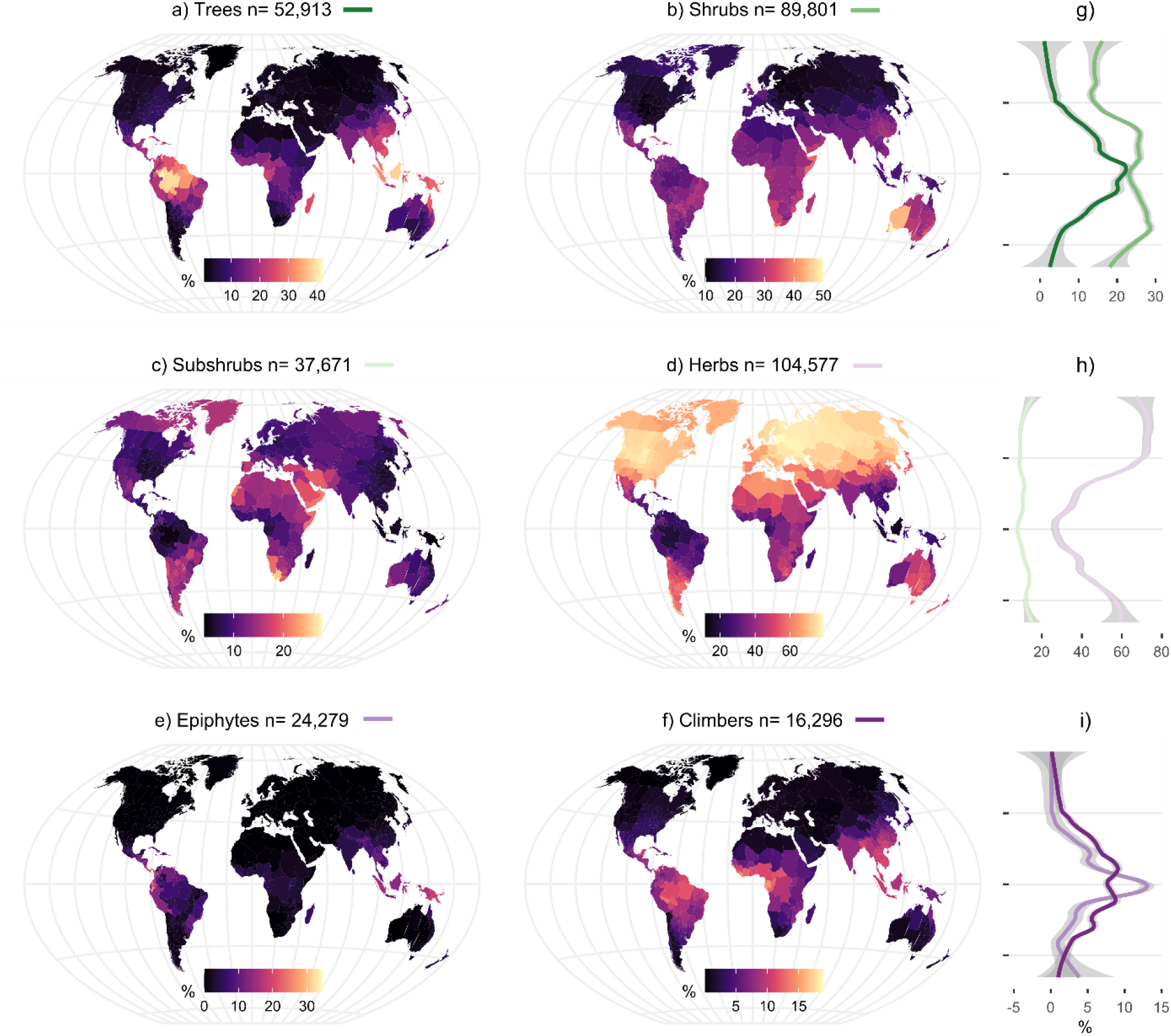
Global patterns of the proportional representation of growth forms (a-f), and their relationship with latitude (panels g-i). Proportional representation was calculated as the number of species within each growth form relative to the combined richness of all growth forms. Note that herbs = terrestrial herbs. Maps are projected following the Winkel tripel projection (Winkel III). See Fig. S1 for equivalent species richness maps.

**Figure 3.**
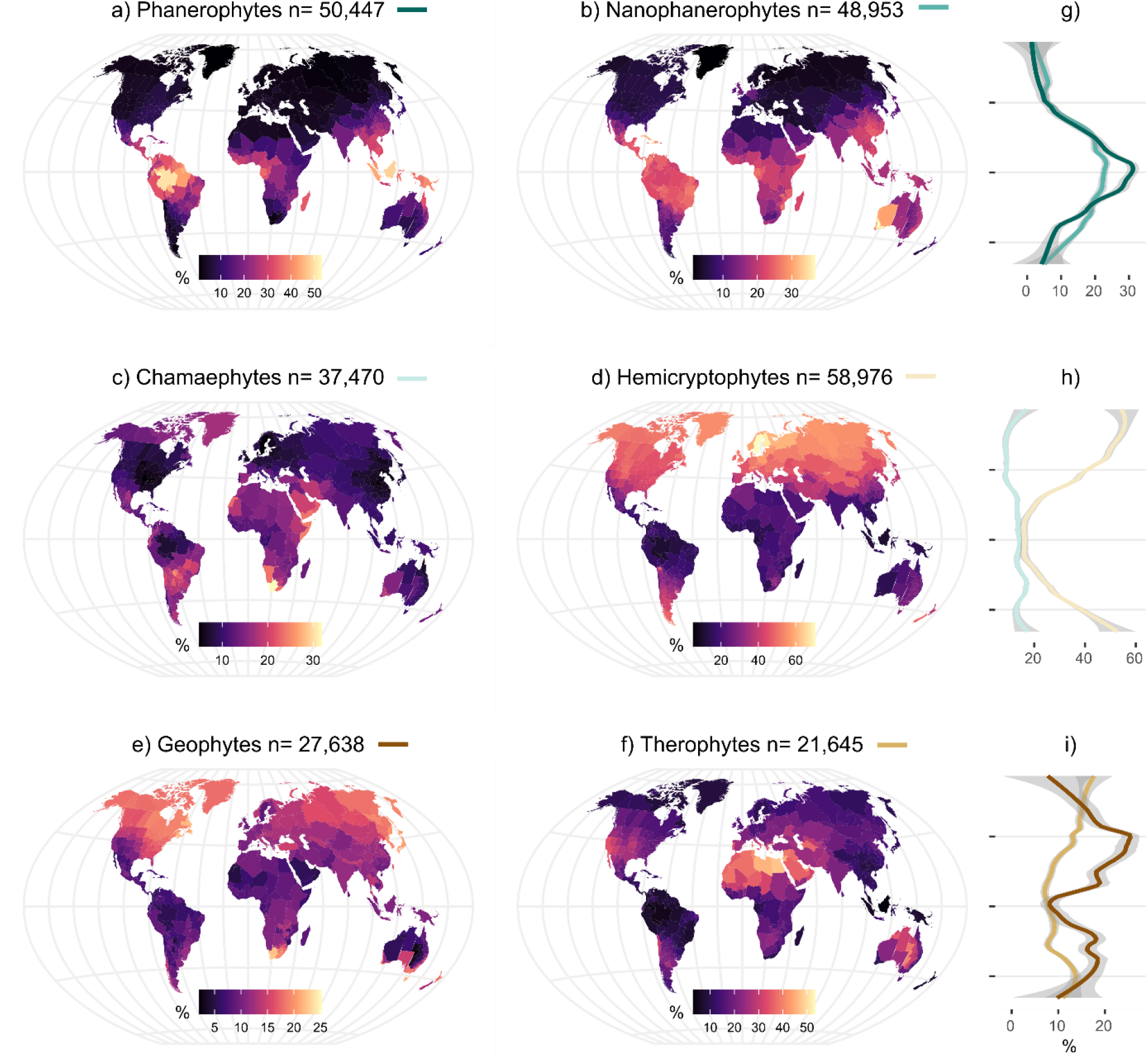
Global patterns of the proportional representation of life forms (a-f), and their relationship with latitude (panels g-i). Proportional representation was calculated as the number of species within each life form relative to the combined richness of all life forms. Note that herbs = terrestrial herbs. Maps are projected following the Winkel tripel projection (Winkel III). See Fig. S2 for equivalent species richness maps.

**Figure 4.**
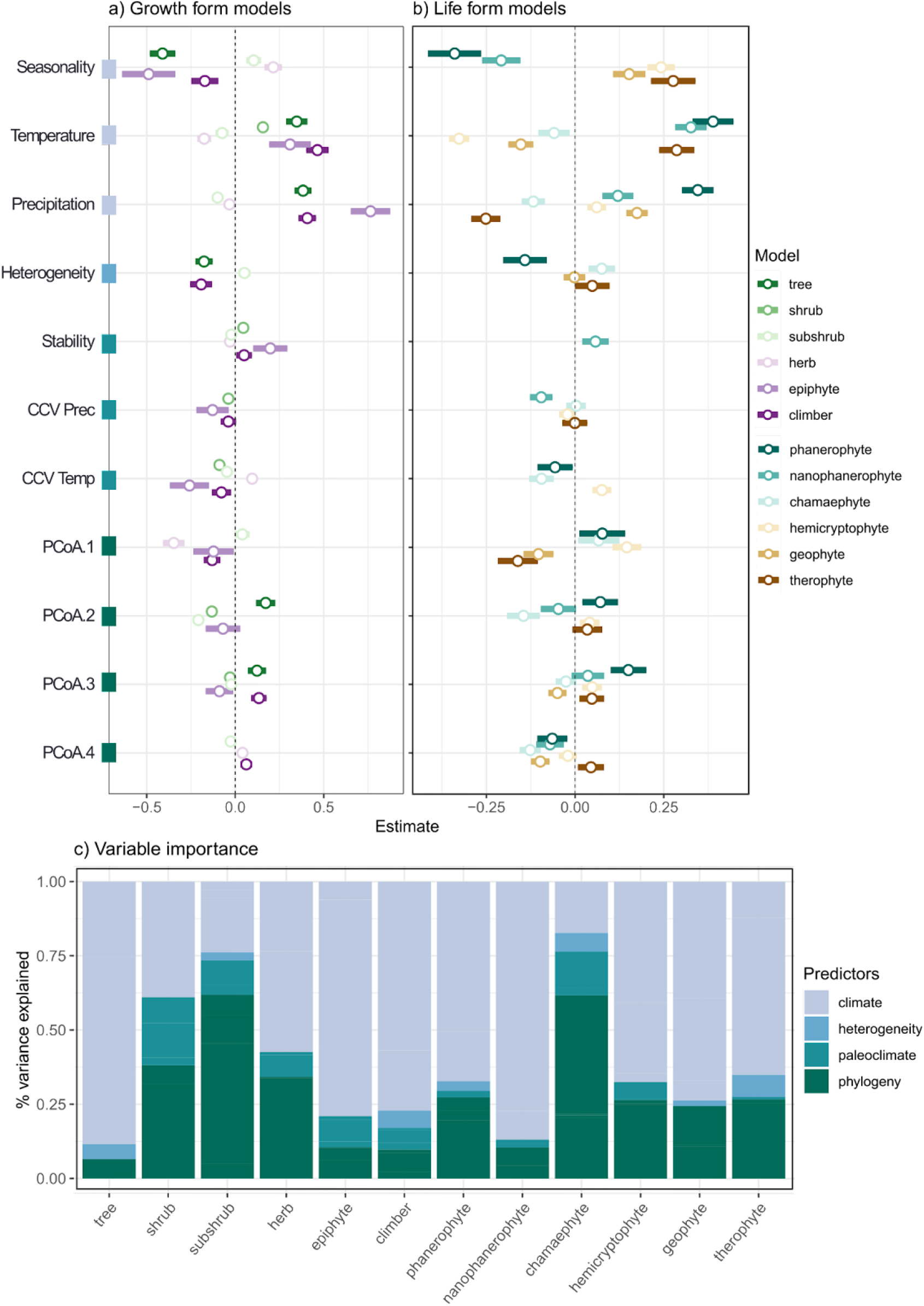
Coefficient plots showing the standardised effects of temperature seasonality (Seasonality, coefficient of variation of temperature), mean annual temperature (Temperature, °C), mean annual precipitation (Precipitation, mm), climate stability (Stability, no units), past climate change velocity in precipitation (CCV Prec, m/a) and temperature (CCV Temp, m/a), climate heterogeneity (Heterogeneity, no units), and phylogeny (PCoA 1-4, as components from a Principal Coordinates Analysis) on the proportional representation of plant (a) growth forms and (b) life forms. Note that the direction of the PCoA axes is not important. Only predictors that made it to the most parsimonious models for each plant form are included. Variable importance (c) plots depict the combined importance of contemporary climate, paleoclimate, environmental heterogeneity, and phylogeny on plant form contributions to global gradients of vascular plant diversity. Detailed model results and data used to create these figures can be found in Table S3, S4.

Despite accounting for only 7% of growth form richness (total number of individuals assigned as growth forms, not species richness), epiphytes comprised up to 35% of the flora of Ecuador, even proportionately representing more than their host trees in 16 regions, and exceeding all other growth forms in two regions (Table S3). The higher contribution of epiphytes to diversity in tropical forests can be related to mean annual precipitation (Est. = 0.76, p = <0.01), and low temperature seasonality, given the strong, inverse relationship between epiphytes and this predictor (Est. = −0.49, p = <0.01). Further corroborating that epiphytes prefer stable climates was their negative association to past climate change velocity (CCV) in temperature (Est. = −0.26, p = <0.01), CCV in precipitation (Est. = −0.13, p = 0.01), and positive association to climate stability (Est. = 0.20, p = <0.01), which only weakly predicted most other growth forms. Climbing plants, which like epiphytes are structurally dependant on trees, displayed strong, positive relationships with precipitation (Est. = 0.41, p = <0.01) and temperature (Est. = 0.46, p = <0.01), and were most prominent in Afrotropical and Neotropical forests (up to 32%). Climbing plants along with trees decreased in proportional representation with increasing climate heterogeneity, highlighting their tight association to tropical forests. Finally, our variable importance analysis revealed phylogeny to be a poor predictor of tree, epiphyte, and climbing growth forms (6-11%), a moderate to strong predictor of herbs and shrubs (34-38%), and a strong predictor of subshrub (62%) representation (Fig. 4b). Together, these predictors explained between 54% and 75% of model variance in our GLMM (marginal R^2^).

### Life forms

As with growth forms, the distribution of life forms varied considerably among the major climate gradients spanning tropical, temperate, and arid ecosystems (Fig. 3). Phanerophytes mirrored the representation of trees and reached their greatest representation in the Neotropics, Afrotropics, and Southeast Asia (up to 53%). This pattern was best predicted by temperature (Est. = 0.39, p = <0.01, Fig. 4a) followed by precipitation (Est. = 0.35, p = <0.01). The proportional richness of nanophanerophytes peaked in Mediterranean forests, but also subtropical dry forests and grasslands (up to 37%, Table S3) and was most strongly related to temperature (Est. = 0.33, p = <0.01). Chamaephytes peaked in representation in regions characterised by deserts and shrublands, tropical grasslands and Mediterranean climates (up to 32%), which was driven by a combined effect of precipitation and phylogeny (Fig. 4a, Table S4). Geophytes, hemicryptophytes, and therophytes displayed opposing patterns of representation, with geophytes being most prevalent in dry Mediterranean (e.g., Cape Province, South Africa), hemicryptophytes in boreal regions (up to 71%), and therophytes in regions with expansive deserts (up to 54%). Despite these geographical differences, the representation of geophytes, hemicryptophytes, and therophytes was strongly related to temperature, with geophytes and hemicryptophytes favouring cold temperate climates, and therophytes favouring hot, seasonal, and dry climates (Fig. 4a, Table S4).

Climate heterogeneity, stability, and past CCV were generally weak predictors of life form representation, accounting for <15% of total variation explained in our variable importance analysis, compared to 65-87% explained by temperature, seasonality, and precipitation for all life forms except chamaephytes for which contemporary climate only explained 17% of variation. Phylogeny, however, while generally a poor to moderate predictor of life form representation (10-27%), was the most important predictor of chamaephyte representation (62%, Fig. 4b). Together, these predictors explained a high degree of model variance, with marginal R^2^ values ranging from 39%-86% (Table S4).

## Discussion

The question of how biodiversity is distributed in space and time is an important and complex topic in ecology and evolutionary biology and requires an integrative perspective on species distributions, ecologically relevant traits, and phylogenetic relatedness. While the knowledge of the distribution of vascular plants has substantially increased with recent advancements in the availability of global datasets (Enquist *et al.*, 2019; Weigelt *et al.*, 2020; Cai *et al.*, 2022), studies on the functional component of plant biodiversity are lagging behind. We quantified the relative contribution of life and growth forms in different regions around the world and found marked differences in their contributions to global gradients of vascular plant diversity. In support of *H_1_*, we found a major shift in the representation by woody perennials in tropical latitudes to almost entirely terrestrial herb-dominated floras in temperate and boreal regions. Notably, while trees may occupy over 80% of biomass in (northern) Europe, America, and temperate Asia (Woodward *et al.*, 2004), we show that they only account for <3% of the vascular plant richness. Terrestrial herbs, in contrast, constitute up to 90% of the vascular flora in these same regions.

Several mechanisms may enable herbaceous species to withstand extreme climate variations. (Zanne *et al.*, 2014). Geophytes, for instance, can survive environmental pressures by utilising belowground storage organs and keeping their perennating buds below the soil surface (Raunkiaer, 1934). In cold climates with frequent frost, hemicryptophytes may shed aboveground biomass, thus insulating their perennating buds located at or just beneath the soil surface (Raunkiaer, 1934; Lubbe *et al.*, 2021). Finally, short-lived therophytes can persist as seeds that lay dormant in seed banks until favourable conditions arise for germination and regrowth (Cross *et al.*, 2015). Although woody perennials typically lack these adaptations, our study revealed that shrubs and subshrubs frequently coexist with herbs in high proportions, suggesting the presence of additional strategies that enable these plant forms to thrive in extreme climates.

Shrubs, for example, are composed of numerous stems that may favour their persistence in unfavourable climates, as injury to one or multiple stems is less likely to result in plant mortality (Götmark *et al.*, 2016). Likewise, the sizable allocation of belowground organs of subshrubs allows individuals to resprout and recover following major disturbances (Giroldo *et al.*, 2017), which could explain their high occurrence in regions characterized by fire and drought (e.g., Australia). Sclerophyllous leaves or succulent stems may provide additional protection to shrubby growth forms in hot, arid environments through the reduction of water loss and increased water storage (Guerra & Scremin-Dias, 2018). These special adaptations to withstand climate extremes make shrubs and subshrubs important contributors of biodiversity in challenging environments where they likely also provide a variety of ecological functions (e.g., habitat, water regulation, soil stability, McKell, 1975).

Trees, on the other hand, reached their greatest proportional representation in tropical forests, which is a well-known pattern from several empirical studies (Keil & Chase, 2019; Cazzolla Gatti *et al.*, 2022). Nevertheless, our findings indicate that the proportion of trees in tropical forests does not match the prevalence of herbs in temperate regions when viewed from a global perspective, corroborating previous studies that a significant component of biodiversity in tropical forests actually comes from non-trees (Gentry & Dodson, 1987; Spicer *et al.*, 2020). Indeed, tropical forests are multi-tiered with epiphytes, climbers, terrestrial herbs, and shrubs also forming important functional components. Epiphytes, for example, constitute just 9.6% of the world’s vascular flora (Zotz *et al.*, 2021), yet can represent up to 39% of regional Neotropical diversity (Taylor *et al.*, 2022), and 50% of local diversity (Kelly *et al.*, 2004). The increase in proportional representation of epiphytes with decreasing scale highlights the importance of scale dependencies in biodiversity analyses, not only due to variation in the contribution by different species, but also the mechanisms underlying these relationships (Crawley & Harral, 2001). Despite the large scale of our analysis, we found in 16 departments spanning Ecuador, Colombia, and Bolivia, epiphytes proportionately exceeded that of their host trees, and in two regions exceeded that of all other growth forms. Our analysis suggests that by neglecting the contribution by non-trees, we are failing to predict half or more of all plant biodiversity in tropical forests.

Although the relative contribution by different plant forms is strongly associated with climate, we show that evolutionary history also plays an important role. The strong effect of phylogenetic relatedness on the representation of subshrubs and chamaephytes may be explained by few plant lineages having the capacity to diversify into the harsh environments typically occupied by these plant forms. Significant phylogenetic conservatism has been found in previous studies on shrubs and subshrubs in arid environments (Zheng *et al.*, 2019), complementing our results that subshrubs are composed of closely related species. Phylogeny may also be important in regions where plant forms are composed of only a few, species-rich genera. One example is of the genus *Aeonium* (Crassulaceae), most of which are succulent chamaephytes endemic to the Canary Islands that have diversified into arid, steep, and rocky habitats inaccessible to most other plant lineages, making them the most successful and specious plant genus across the archipelago (Jorgensen & Olesen, 2001). While we did not include oceanic islands in this study, future investigations on which plant forms are most prone to diversifying on islands may bring us closer to understanding why some plant lineages have more species than others. Conversely, phylogeny had only a weak effect on the proportional representation of epiphytes, despite ~70% of all epiphytes being orchids (Zotz *et al.*, 2021). One possible reason for this is that epiphytes are so strongly coupled to atmospheric conditions, that this may dilute any phylogenetic signal. Our results support this theory as in most cases, epiphytes displayed stronger relationships with climate than any other plant form (Fig. 4a).

Past climate change left only a small legacy on present day plant form distributions, corroborating our *H_2_* that life and growth form contributions are primarily explained by contemporary climate. Terrestrial herbs, for example, were the only plant form positively associated with rapid changes in temperature, demonstrating a major advantage of herbs to exploit disturbed habitats and persist through major climate changes. Global shifts in rainfall regimes are already changing the proportional representation of plant forms in some biomes such as tropical savannahs (Zhang *et al.*, 2019), which is not reflected in species richness alone. This reaffirms the importance of studying plant forms alongside species richness to fully understand the distribution of biodiversity.

## Conclusion

This study highlights variation in the importance of different plant forms to diversity gradients worldwide, providing a much-needed quantification for long-standing ideas and concepts structuring plant assemblages. In line with our expectations, we confirm that the relative contribution of plant forms to global gradients of vascular plant diversity is strongly related to life and growth form dependent responses of plants to regional environmental conditions. Temperature and precipitation continue to be key predictors of plant biodiversity, although we also show that phylogeny matters, particularly for plant forms occupying niche habitats or harsh climates (e.g., subshrubs, chamaephytes). Finally, we highlight the usefulness of considering life and growth forms in biodiversity analyses as a standardised comparison of floras among regions, which in the future may shed light on the evolutionary pressures constraining plant trait distributions or increase our understanding of how different vegetation types may respond to global climate change.

## Author contributions

AT, PW, PD, LC, HK conceived the study, AT collated the data from WCVP, PW and HK created the GIFT database with recent input by PD, AT ran the analyses which were discussed with PW, AT wrote the draft, and all authors significantly contributed to revisions.

## Data availability statement

All aggregated data used for analyses is included in supporting information Table S3.

